# Robust Input Disentanglement Through Dendritic Calcium-Mediated Action Potentials

**DOI:** 10.1101/2025.06.10.658823

**Authors:** Sima Hashemi, Shirin Shafiee, Christian Tetzlaff

**Affiliations:** III. Institute of Physics – Biophysics, Faculty of Physics, University of Göttingen, Göttingen, Germany; Group of Computational Synaptic Physiology, Department of Neuro- and Sensory Physiology, University Medical Center Göttingen, Göttingen, Germany

## Abstract

In daily life, living beings encounter a continuous stream of mixed information, which has to be disentangled by the brain to form proper representations. Using computational modeling, we demonstrate that the interplay between dendritic calcium-mediated action potentials (dCaAPs) with synaptic plasticity and rewiring can enable single neurons to successfully perform this complex task. Compared to other types of dendritic spikes, dCaAPs exhibit a high triggering threshold, large, but graded spike amplitude, with lower amplitudes for stronger synaptic inputs. We show that these properties enable neurons to successfully learn to represent discrete items from a continuous input stream by facilitating the clustering of synapses with temporally correlated presynaptic activities onto the same dendritic branch. In comparison to NMDA spikes, dendrites generating dCaAPs can form representations of individual items more efficiently, independent of the temporal order of their presentation during learning — whether randomly, sequentially, as part of a random stream of simultaneously shown input items, or even as items with shared properties. Thus, our results provide further evidence about the critical role of dCaAPs for the computational capabilities of single neurons.

## Introduction

Early theoretical frameworks in neuroscience often modeled neurons as *point neurons*, simplifying them to entities that integrate incoming signals and produce outputs without accounting for the intricate morphology of dendrites and axons [38, 46, 48]. This simplification was initially deemed acceptable due to its mathematical convenience and the limitations of experimental techniques available at the time [48]. However, it later became evident that such simplification overlooked critical aspects of neuronal computation, as described by cable theory, which mathematically characterizes how electrical signals attenuate and integrate along dendrites [43]. Nevertheless, the active role of dendrites in computational processes had long been overlooked [27], until the advent of advanced, high-resolution recording and imaging techniques revealed the ability of dendrites to generate localized electrical events—known as dendritic action potentials [2, 10, 11, 18]. This discovery challenged the conventional point neuron model by demonstrating that individual neurons, through their dendritic structures, can execute computations once believed to require entire neural networks [29, 30, 54, 55].

Recent studies have highlighted the crucial role of dendritic computations in sensory perception, learning, and memory formation in living animals, further underscoring their functional significance within brain circuits [11, 24, 26, 32, 47, 50]. Experimental studies have shown that the dendritic spikes can be initiated by the coordinated activation of multiple synapses within a specific dendritic segment [31], driving synaptic plasticity, even in the absence of somatic firing [14]. Such localized plasticity mechanisms could facilitate the formation of so-called synaptic clusters that consist of functionally correlated synapses being situated in close proximity onto the same dendritic branch [19, 23, 28]. For instance, dendritic Ca^2+^ spikes in distinct apical tuft branches of layer 5 pyramidal neurons have been shown to correspond to the learning of different motor tasks, with each branch specializing in a specific task [9]. These findings suggest that dendritic spikes enable branch-specific learning rules expanding the computational and learning capabilities of individual neurons [8].

Computational studies have shown that synaptic clustering and dendritic spikes facilitate branch-specific computations and memory compartmentalization [28, 39, 41, 42]. A recent study proposed that the combination of dendritic spikes with local plasticity suggests a mechanism for the emergence of stable synaptic clusters formed by temporally correlated inputs onto dendritic branches [28]. The authors suggest that this mechanism is enabled by NMDA (N-methyl-D-aspartate) spikes, which act as a form of dendritic nonlinearity [28]. NMDA spikes are one of three known types of dendritic spikes, alongside sodium and calcium spikes [4, 20]—each characterized by distinct electrophysiological properties, including activation thresholds, durations, and amplitudes, as well as unique functional roles. Among these, NMDA and calcium spikes are well-characterized, branch-specific regenerative events that occur when several nearby synapses are activated nearly simultaneously [3, 16]. Unlike sodium spikes, which typically propagate along the axon and soma, NMDA and calcium spikes are often confined to small dendritic segments, making them ideal for local synaptic integration and clustering [3, 4, 5, 6, 16]. It remains to be explored how the synaptic clustering capacity of dendrites depends on the specific type of dendritic spike, and how NMDA and calcium spikes differ in their influence on synaptic organization.

Dendritic calcium-mediated action potentials (dCaAPs) are a newly discovered subclass of calcium spikes that exhibit unique properties: they are triggered at high input thresholds and display a non-monotonic amplitude response, peaking at threshold and attenuating with stronger stimuli [12, 33]. Computational studies have shown that the unique properties of dCaAPs enable them to support the classification of linearly non-separable inputs within individual dendritic branches [12]. However, the influence of these dendritic spikes on synaptic clustering and the disentanglement of input streams remains unknown.

In this study, we examine how the unique properties of dCaAPs influence dendritic computation and compare their role in synaptic cluster formation to that of NMDA-mediated dendritic spikes. To achieve this, we model a compartmental neuron with active dendritic compartments, incorporating both structural and functional plasticity mechanisms. Our results demonstrate that dendrites equipped with dCaAPs outperform those with NMDA spikes in the formation of robust synaptic clusters across various input protocols. These findings emphasize the unique computational contributions of dCaAPs and highlight their potential role in facilitating input stream disentanglement, thereby enhancing the overall information-processing capacity of individual neurons.

## Materials and Methods

The model used in this study builds upon the framework introduced by Limbacher et al. (2020) [28] that considered a multi-compartment neuron with NMDA-based dendritic spikes in conjunction with plasticity mechanisms. In addition, we integrate the dCaAP model proposed by Gidon et al. (2020) [12], as model for dendritic spike generation. In the following, we describe the key components drawn from Limbacher et al. and detail our modifications to the dendritic nonlinearity, along with a comparison of our results to those of the original framework. The code used for simulations and analyses in this study is available at: https://github.com/simahashemi/Single-Cell-Synaptic-Clustering.

### Neuron and dendrite model

The compartmental model of a neuron comprises a soma connected to 12 dendritic branches (or compartments), each functioning as a leaky integrator. The dynamics of the somatic membrane potential are described by the following differential equation:

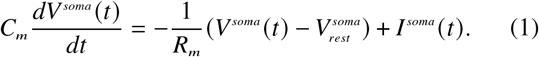

The term *I*^*soma*^ represents the forward current from the dendrites to the soma with membrane resistance *R*_*m*_, membrane capacity *C*_*m*_ and resting potential 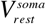 (see Table 1).

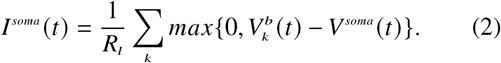

**Table 1.** Model parameters.

| Parameter | Description | Value | Reference | Parameter | Description | Value | Reference |
| --- | --- | --- | --- | --- | --- | --- | --- |
| $C_m$ | Somatic membrane capacity | $250pF$ | [28] | $\tau^{amp}$ | dCaAP maximum amplitude constant | 0.39 | [12] |
| $R_m$ | Somatic membrane resistance | $40M\Omega$ | [28] | $\tau^{rise}$ | dCaAP current rise time constant | $3ms$ | [12] |
| $V_{soma}^{rest}$ | Somatic rest potential | $-70mV$ | [28] | $\tau^{decay}$ | dCaAP current decay time constant | $0.4ms$ | [12] |
| $R_l$ | Longitudinal resistance | $2\Omega$ | [28] | $\Delta t^s$ | dCaAP current time delay | $21ms$ | [12] |
| $\rho$ | Firing rate scale | $2.5Hz$ | [28] | $V_{rest}$ | Dendritic rest potential | $-70mV$ | [28] |
| $\beta$ | Firing rate constant | 0.5 | [28] | $\eta$ | Learning rate | 0.002 | [28] |
| $V_{soma}^{thr}$ | Somatic firing threshold | $-55mV$ | [28] | $C_{str}$ | Scale factor in structural plasticity | 5.5 | [28] |
| $C_b$ | Dendritic membrane capacity | $250pF$ | [28] | $\lambda$ | Scale factor in structural plasticity | 10 | [28] |
| $R_b$ | Dendritic membrane resistance | $40M\Omega$ | [28] | $C_w$ | Scale factor in structural plasticity | $0.55nA^{-1}$ | [28] |
| $V_{rest}^b$ | Dendritic rest potential | $-70mV$ | [28] | $c_\theta$ | Weight scale factor | $1nA$ | [28] |
| $\tau_{syn}$ | Synaptic time constant | $2ms$ | [28] | $N_{syn}$ | Constraint on the number of synapses | 20 | [28] |
| $c_{ds}$ | Scale factor for plateau duration | $0.04s^2/mV$ | [28] | $C_{func}$ | Scale factor in functional plasticity | 1.5 | [28] |
| $V_s$ | Initial amplitude of sodium spikelet | $5mV$ | [28] | $\gamma$ | Scale factor in functional plasticity | 0.2 | [28] |
| $\tau_s$ | Time constant of sodium spikelet | $4ms$ | [28] | $\tau_x$ | Time constant of presynaptic trace | $20ms$ | [28] |
| $V_{ds}$ | Amplitude of plateau potential | $-30mV$ | [28] | $C_{STDP}$ | Scale factor in STDP plasticity | 3.2 | [28] |
| $A_k^{max}$ | dCaAP maximum amplitude | $43.8mV$ | [12] | $V_{thr}^{STDP}$ | Threshold for STDP plasticity | $-67mV$ | [28] |
| $V_{thr}^{dCaAP}$ | dCaAP threshold | $-36mV$ | [12] | $C_{noise}$ | Scale factor in noise term | 0.035 | [28] |

Here, *R*_*l*_ represents the longitudinal resistance and 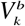 denotes the membrane potential of dendritic branch *k*. When a positive gradient in somatic voltage is present, the firing rate of the soma will be modulated according to the following equation with firing rate scale *ρ*, firing rate constant *β*, and somatic firing threshold 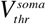:

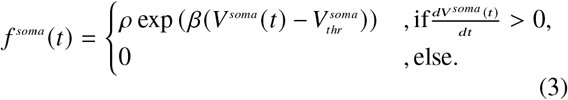

Upon firing, the soma enters a refractory period of 5 ms, during which the somatic potential is reset to 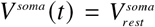.

The membrane potential of dendritic branch *k*, denoted as 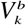, is characterized by a membrane resistance *R*_b_, a membrane capacitance *C*_*b*_ and resting potential 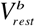 (see Table 1), and is calculated as follows:

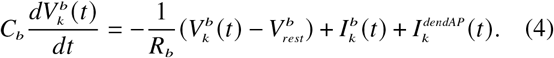

The input current to each dendritic branch *I*_*b*_ is calculated as the weighted sum of the alpha functions corresponding to the spikes of the input neurons: 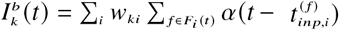.

Here, *w*_*ki*_ is the weight of the connection between dendritic branch *k* and input neuron *i*. The *F*_*i*_ *t* represents the set of all input spikes from neuron *i* occurred before *t*, where each spike is denoted by 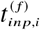. The alpha function presented above represents presynaptic action potentials 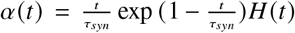 with *τ*_syn_ being the synaptic time constant. Additionally, 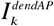 represents the excess current resulting from a dendritic spike on branch *k*. Throughout this study we will consider two different types of dendritic spikes: NMDA-based dendritic spikes and dendritic calcium-mediated action potentials (dCaAPs). The mathematical description of NMDA-based dendritic spikes is based on Limbacher et al. (2020), while the used dCaAP-model is based on Gidon et al. (2020).

#### dCaAP spikes

When the membrane potential of a dendrite exceeds the dCaAP threshold, a dendritic spike is elicited, 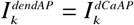, and that branch enters an absolute refractory period as long as 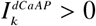. During the refractory period 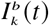 is set to be zero and no new dendritic spike can be triggered. We modeled the dCaAP current with rise and decay time constants (*τ*^*rise*^ and *τ*^*decay*^) as defined by Gidon et al. (2020) [12] such that

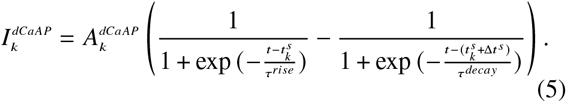

In the absence of dCaAP spikes, this current is zero.

The maximum amplitude for 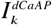 is given as follows:

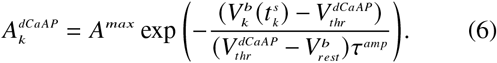

The parameters *A*^*max*^ and 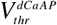 denote the mean peak amplitude of the dCaAPs and the membrane potential threshold for dCaAP initiation, respectively, as derived from experimental measurements [12]. The 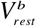 parameter represents the resting membrane potential of the dendrites, while 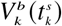 denotes the membrane potential of dendritic branch *k* at the time of the dCaAP spike 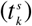.

#### NMDA spikes

The model of the NMDA-based dendritic spike comprises two components : a sodium spikelet with a peak potential of *V*_*s*_ = *−* 25*mV*, followed by a plateau at a constant potential of *V*_*ds*_ = *−* 30*mV* [28]. Different to the model of dCaAPs, instead of providing an additional current *I*^*dendAP*^, the model of the NMDA-based dendritic spikes directly sets the membrane potential of the corresponding dendritic branch:

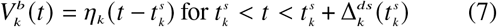

with

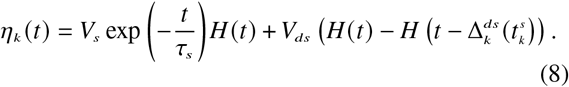

*H*(·) is the Heaviside function. The duration of the plateau is a function of the gradient of the membrane potential at the moment of firing: 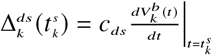.

While the plateau potential is maintained, the dendritic branch is in an absolute refractory state, indicating that no further regenerative events can occur. The parameter values for both the dCaAP and NMDA-based dendritic spike models are listed in Table 1.

### Synaptic and structural plasticity

Considering the biological and structural limitations of dendritic branches, each branch in our model has a limited capacity for the realization and maintenance of connections with input neurons. Therefore, we introduce the variable *θ*_*ki*_, which indicates the state of a potential synapse over time. A positive value of this variable indicates the existence of a functional connection between input neuron *i* and dendritic branch *k*, with a weight defined as *w*_*ki*_ = *c*_*θ*_ *θ*_*ki*_ (*w*_*ki*_ ∈ [0, 8]). Thus, if *θ*_*ki*_ falls below zero, a functional synapse is converted into a potential synapse, and vice versa, thereby establishing dynamic synaptic wiring. The dynamics of synaptic states are governed by a differential equation that consists of several components:

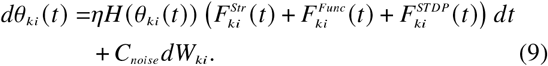

The Heaviside function (*H*(·)) ensures that the following plasticity terms only influence functional synapses (*θ*_*ki*_ *t* > 0).

#### Structural plasticity 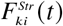

This term describes the synaptic rewiring dynamics, providing a structural constraint that regulates the maximum number of functional synapses a branch can establish. A schematic of this function is plotted in Figure 1A.

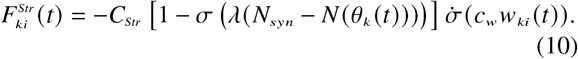

**Figure 1:**
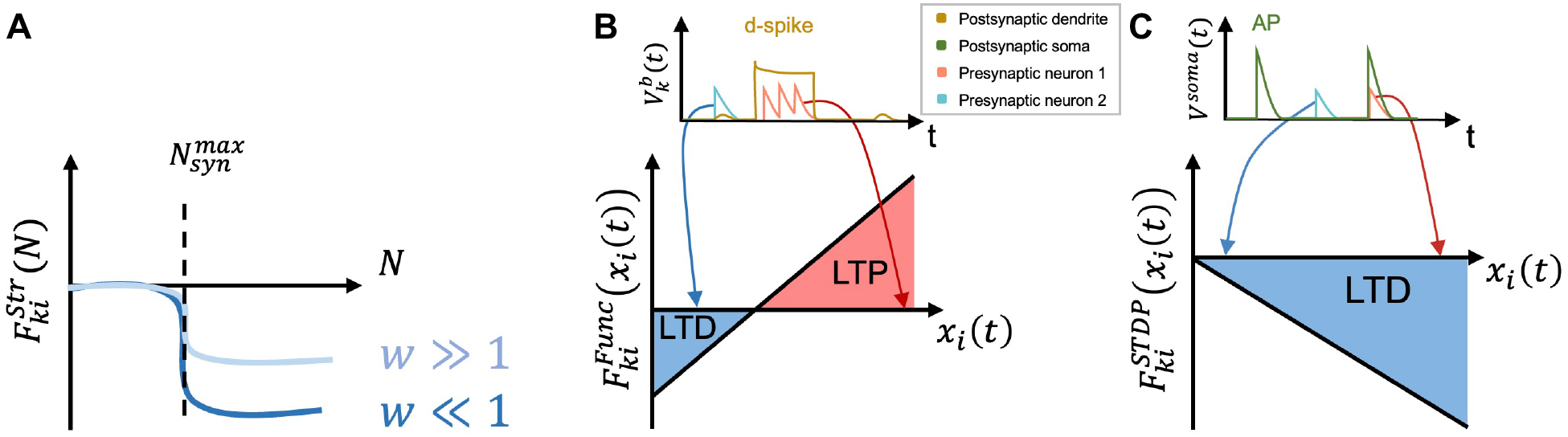
Synaptic and structural plasticity. **A)** *Structural Plasticity*. This plasticity rule enforces a constraint on the dendritic branches, limiting the number of synaptic connections to a maximum threshold 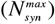. When the number of connections on a branch exceeds this limit, all of the connections on that branch are weakened in proportion to their synaptic weights. As a result, stronger synapses with larger synaptic weights (*w* >> 1) undergo minimal weakening, while weaker synapses with smaller weights (*w* << 1) are more substantially weakened. **B)** *Functional Plasticity*. When a dendritic spike occurs in a branch, the synaptic weights of all connections on that branch are adjusted based on the activity of the presynaptic neurons. Connections associated with presynaptic neurons that were highly active during the dendritic spike and plateau phase (red trace), are strengthened (LTP). Conversely, connections linked to less active presynaptic neurons (blue trace) during this period are weakened (LTD). **C)** *STDP Plasticity*. Once an Action Potential (AP) is generated, the STDP plasticity induces an LTD effect on all the connections on branches with a membrane potential above the threshold 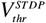. The degree of synaptic weakening depends on the timing of the presynaptic spike relative to the postsynaptic AP. When the presynaptic spike occurs just before or during the AP (red trace), the weakening is more pronounced, whereas a later presynaptic spike (blue trace) leads to a less substantial reduction in synaptic weight.

Here, the parameters *C*_*Str*_, *λ*, and *c*_*w*_ are positive scale factors and constants, as specified in Table 1. The sigmoid term *σ*(·) restricts the number of synapses on each dendritic branch *N θ*_*k*_ to a maximum value of *N*_syn_ (set to 20 in this study). The total number of synapses on each branch is calculated by 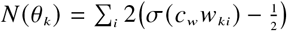. The sigmoid derivative term 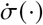 implements a dynamic that preferentially penalizes weaker synapses compared to stronger ones when *N θ*_*k*_ exceeds *N*_syn_. This mechanism is consistent with experimental findings indicating that larger spines (which correspond to stronger synapses) have been found to be more persistent in vivo than smaller spines [13, 15, 35, 52].

#### Functional plasticity 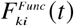

This term describes the processes that potentiate or diminish the synaptic connections based on the activity of the connected input neuron during dendritic spikes (Figure 1B). This functional plasticity mechanism is implemented as follows:

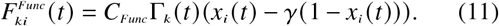

The parameter *C*_*Func*_ represents the scale factor for functional plasticity, and *γ* is a constant that determines the threshold activity at which the transition from potentiation to depression occurs (Table 1). The function Γ_*k*_ *t* acts as a dendritic spike indicator, taking a value equal to one during a spike and zero at all other times. *x*_*i*_ *t* represents the presynaptic activity trace of input neuron *i*: 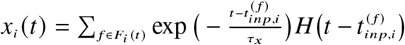. Here, *τ*_*x*_ denotes the time constant for presynaptic traces (Table 1).

#### Spike-timing dependent plasticity (STDP) 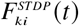

Previous studies have shown that a spike-timing-dependent mechanism results in each somatic spike inducing a transient depression of all synapses that have recently undergone presynaptic spiking events [25]. Consequently, 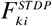 is introduced to induces long-term depression (LTD) on the synapses that were active during the somatic spike, provided that the dendritic branch, where they are located, exceeds a threshold 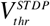 (see Table 1).

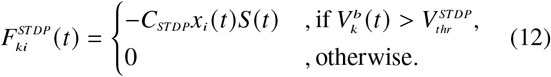

Here *C*_*STDP*_ is a positive scale factor for the STDP term (see Table 1). *S t* represents the postsynaptic somatic spike train: *S* (*t*) = ∑*fδ*(*t*−*t* ^(*f*)^). Figure 1C represent a schematic of this plasticity term.

#### Synaptic noise

Finally, stochasticity is introduced into the model through the last term in Equation 9. In this context, *dW*_*ki*_ represents increments from a standard Wiener process, while the temperature parameter *C*_*noise*_ is a constant that modulates the intensity of this stochastic component.

## Results

Neurons utilize different dendritic spike types to process information, being received from a continuous stream of inputs. To understand the role of dendritic calcium-mediated action potentials (dCaAPs), we model a neuron consisting of a soma and several dendritic branches (Fig. 2). Synapses are plastic, leading to changes in their weights and implementing synaptic rewiring. The learning phase involves the presentation of assemblies to the neuron, utilizing four distinct protocols. Each assembly is composed of 40 input neurons. The learning phase consists of alternating periods of 300 ms activation window and 200 ms resting window. During each activation window, all 40 neurons within a selected assembly fire as a Poisson spiking group at a frequency of 35 Hz, while the remaining input neurons fire randomly at the basal frequency of 1 Hz. In resting windows, all input neurons fire at 1 Hz.

**Figure 2:**
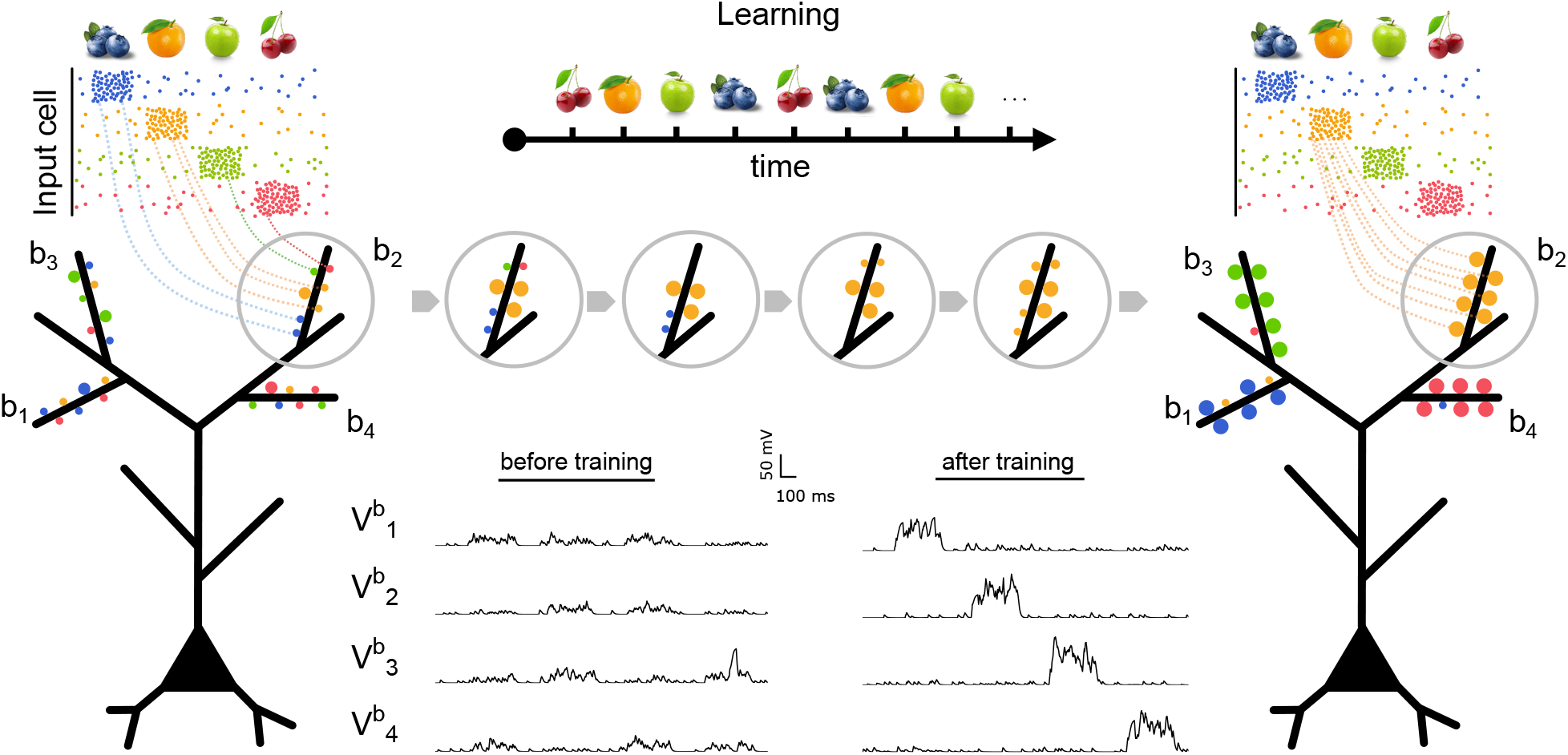
Schematic of learning process. The left panel shows the neuron prior to learning, where input cells are randomly connected to dendritic branches with low synaptic weights. At this stage, the membrane potential of the branches is noisy and does not reflect any response to a specific input assembly. During the learning phase, assemblies are presented to the model according to defined protocols. For clarity, each assembly is represented by a distinct fruit symbol and color. As learning progresses, dendritic spikes and synaptic plasticity mechanisms lead to the weakening or removal of some synapses and the strengthening or formation of others. This process gives rise to synaptic clusters on individual branches. After learning (right panel), each dendritic branch becomes selectively tuned to a specific assembly. A branch is considered to have learned an assembly if it contains more than 10 synapses with a total weight greater than 50 originating from that assembly. In the post-learning state, the membrane potential of each branch shows increased activity during the presentation of its associated assembly, indicating successful specialization.

Before learning, input neurons are randomly connected to the dendritic branches of the neuron via functional connections. Throughout the learning process, synaptic rewiring leads to the pruning of some of the initial random functional connections, while certain potential synapses are activated and transformed into functional synapses. Additionally, the synaptic weights of functional synapses are modified. Figure 2 illustrates a schematic of the neuron model before and after learning. Each assembly encodes an input item like a fruit, being presented to the neuron during the learning phase. The resulting synaptic adaptations lead to a clustering of synapses receiving input from the same assembly, onto the same dendritic branch. By this, the dendritic branch shows the maximum amplitude of its membrane potential in response to the presentation of the assembly that formed the synaptic cluster. In other words, after the learning phase, a dendritic branch hosting one synaptic cluster, is tuned towards the related assembly - the dendritic branch became a “learned branch”. We define the efficiency of a neuron after learning as its ability to represent and detect the maximum number of assemblies while engaging the fewest number of branches, thereby conserving resources and increasing storage capacity. To quantify this efficiency for different dendritic spike types (here dCaAP and NMDA-based), we introduce an efficiency factor *ξ*:

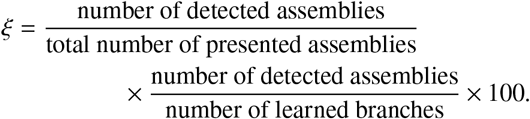

The efficiency factor is *ξ* = 0% if no input assembly is represented or detected by any dendritic branch of the neuron. Conversely, if the neuron detects all *N* presented assemblies using *N* distinct dendritic branches, the efficiency reaches *ξ* = 100% (see Figure S1 in the Supplementary Materials).

To examine the efficiency of the neuron in different contexts, we explored four distinct learning paradigms: random presentation of input assemblies, sequential presentation, random presentation of overlapping assemblies, and the co-active presentation of assemblies. The following subsections describe each of these protocols along with their corresponding results. For a more detailed examination of each protocol, we refer the reader to the Supplementary Materials.

### Random inputs

In the first protocol, we explored the influence of NMDA- and dCaAP-generated dendritic spikes on a neuron’s ability to learn and represent individual assemblies from a continuous input stream, which presents one assembly at a time in a random order. To this end, during the learning phase, one assembly was randomly selected from the eight available assemblies and activated as a Poisson spiking group at a frequency of 35 Hz during each activation window, followed by a resting window. This process was iteratively repeated over 1000 seconds of learning. Afterward, the model was tested to assess its capacity to encode and retain disjoint assemblies from randomized inputs (see Figures S2 and S3 in the Supplementary Materials). The results from 10 simulation runs for both neuron models—one with NMDA-triggered and the other with dCaAP-triggered dendritic spikes—are shown on the left side of Figure 3. The NMDA model detects an average of 5.1 ± 1.1 assemblies with 8.7 ± 1.5 learned branches, while the dCaAP model detects an average of 5.7 ± 1.1 assemblies and exhibits 9.5 ± 2.1 learned branches. Consequently, both models demonstrate similar efficiencies, with average efficiency values of 38.8% ± 14.9% for the NMDA model and 43.7% ± 11.0% for the dCaAP model.

**Figure 3:**
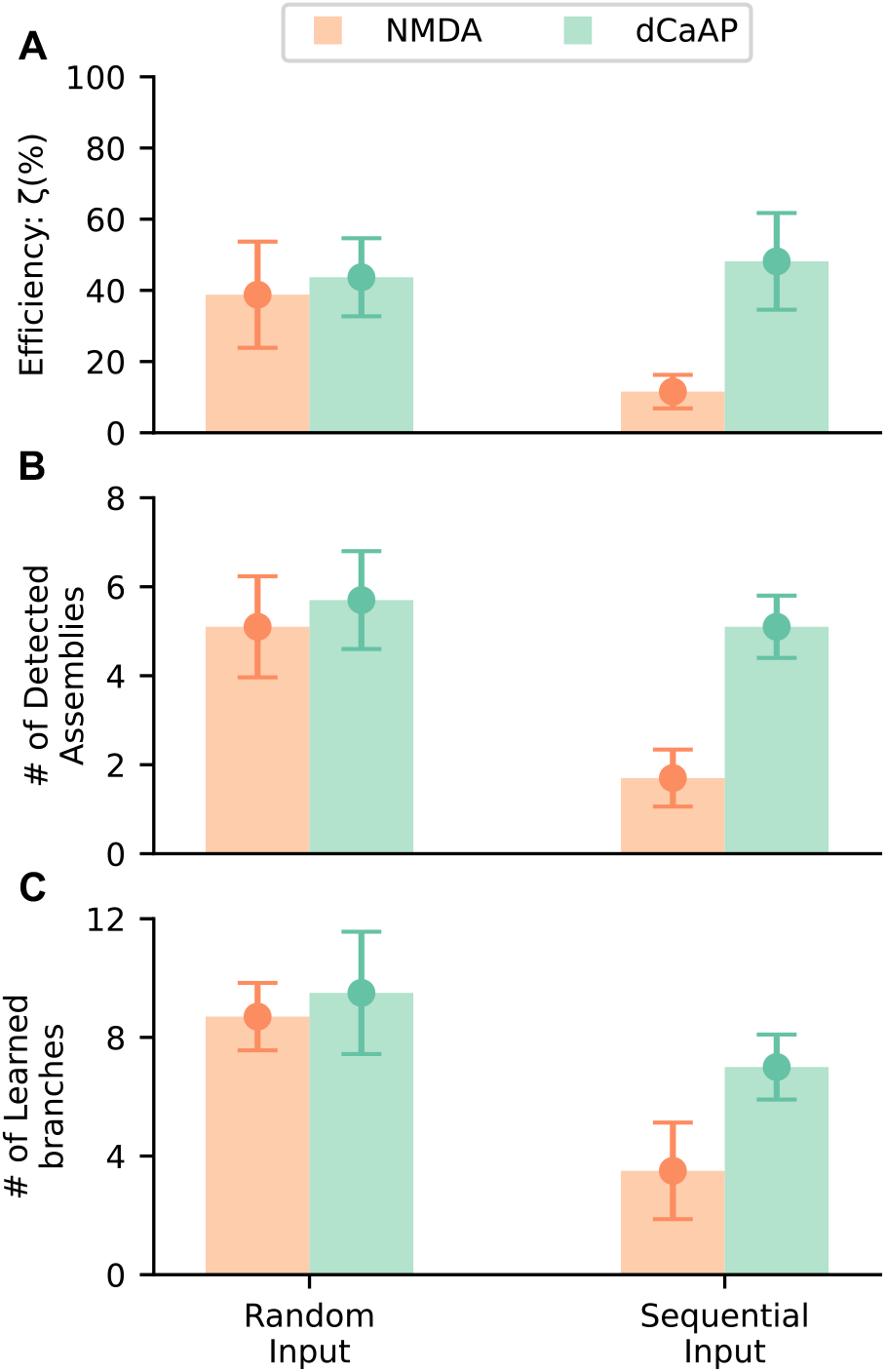
Random vs. sequential input protocols. Simulation results from 10 trials under random and sequential input protocols are shown for a neuron model with NMDA-based dendritic spikes (orange) and a model with dCaAPs (green). **A)** average efficiencies; **B)** number of detected assemblies; **C)** number of learned branches.

### Sequential inputs

The brain is constantly exposed to a stream of events, yet it is able to learn from them without erasing previously acquired memories. To replicate this process, we applied a sequential learning protocol to observe whether the NMDA- and dCaAP models could retain previously learned assemblies while incorporating new ones, without overwriting earlier memories. To test this, we divided the learning phase into eight intervals of 125 seconds each. One assembly was activated at 35 Hz rate only during one learning interval, while the remaining assemblies fired at 1 Hz, resulting in a sequential presentation of assemblies. The results of this experiment are presented on the right side of Figure 3. The NMDA-based neuron model detected an average of 1.7 ± 0.6 assemblies using 3.5 ± 1.6 learned branches, yielding an efficiency of 11.5% ± 4.7%. This efficiency is significantly lower than the efficiency observed in the random input paradigm. In contrast, the dCaAP model detected on average 5.1 ± 0.7 assemblies with 7.0 ± 1.1 learned branches, resulting in an efficiency of 48.1% ± 13.6%, which is comparable to this model’s efficiency for the random input protocol. These findings suggest that, unlike the NMDA model, the dCaAP model exhibits resilience to the patterns of input presentation during the learning process. Additionally, the weight traces of both models during the learning phase are shown in Figure 4. These plots indicate that the NMDA model experiences significant overwriting of earlier learned information, leading to instability in synaptic weights as new inputs are presented. In contrast, the dCaAP model demonstrates robust synaptic stability, preserving previously acquired weights even in the presence of subsequent inputs. On the right side of this figure, the post-learning states of the neurons are illustrated for both models. In alignment with the synaptic weight trends shown on the left, the NMDA model predominantly allocates most of the branches to the last presented input (represented in red). Conversely, the dCaAP model exhibits clear branch-specific specialization, where each dendritic branch reliably encodes a distinct input assembly. These observations highlight the dCaAP model’s enhanced capacity for preserving learned representations and its resilience to overwriting, effectively mitigating forgetting.

**Figure 4:**
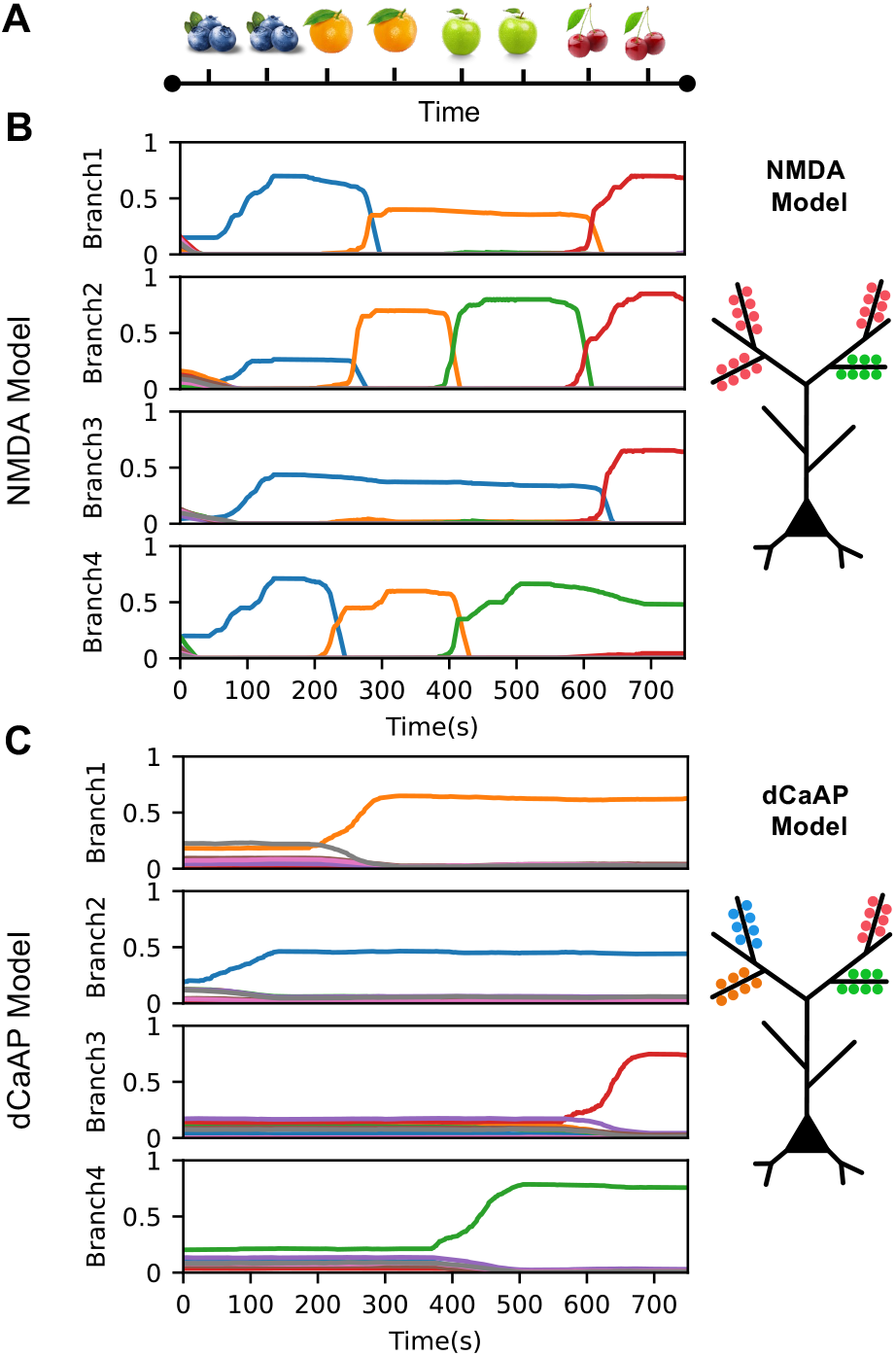
Synaptic dynamics in response to sequential inputs. **A)** Schematic of the sequential learning protocol, where all assemblies (depicted as distinct fruits) are presented consecutively, each activated for 125 seconds. **B)** Synaptic weight dynamics across four dendritic branches of the NMDA-based neuron model. Weights are normalized such that the maximum possible sum for any assembly is 1. The NMDA model shows substantial overwriting, with later inputs dominating earlier ones. **C)** As in B, but for the dCaAP model. Synaptic weights exhibit long-term stability despite exposure to novel inputs.

### Overlapping inputs

Up to this point, all input protocols involved completely disjoint assemblies. However, real-world stimuli often share features resulting in overlapping assemblies. For instance, the similarity in color of a cherry and a red apple results in overlapping activation of their corresponding assemblies (Figure 5A). In this protocol, we presented the assemblies to the models in random order, with varying degrees of overlap — 25%, 50%, and 75% — and evaluated the overlap’s impact on learning outcomes. For instance, in the case of 50% overlap, half of the assembly population is shared with other assemblies and becomes active when those assemblies are activated. The results are presented in Figure 5B. While the performance of the NMDA and dCaAP models is similar in the case of no overlap, the dCaAP model outperforms the NMDA model when learning with highly overlapping assemblies (see also Figure S4 in the Supplementary Materials).

**Figure 5:**
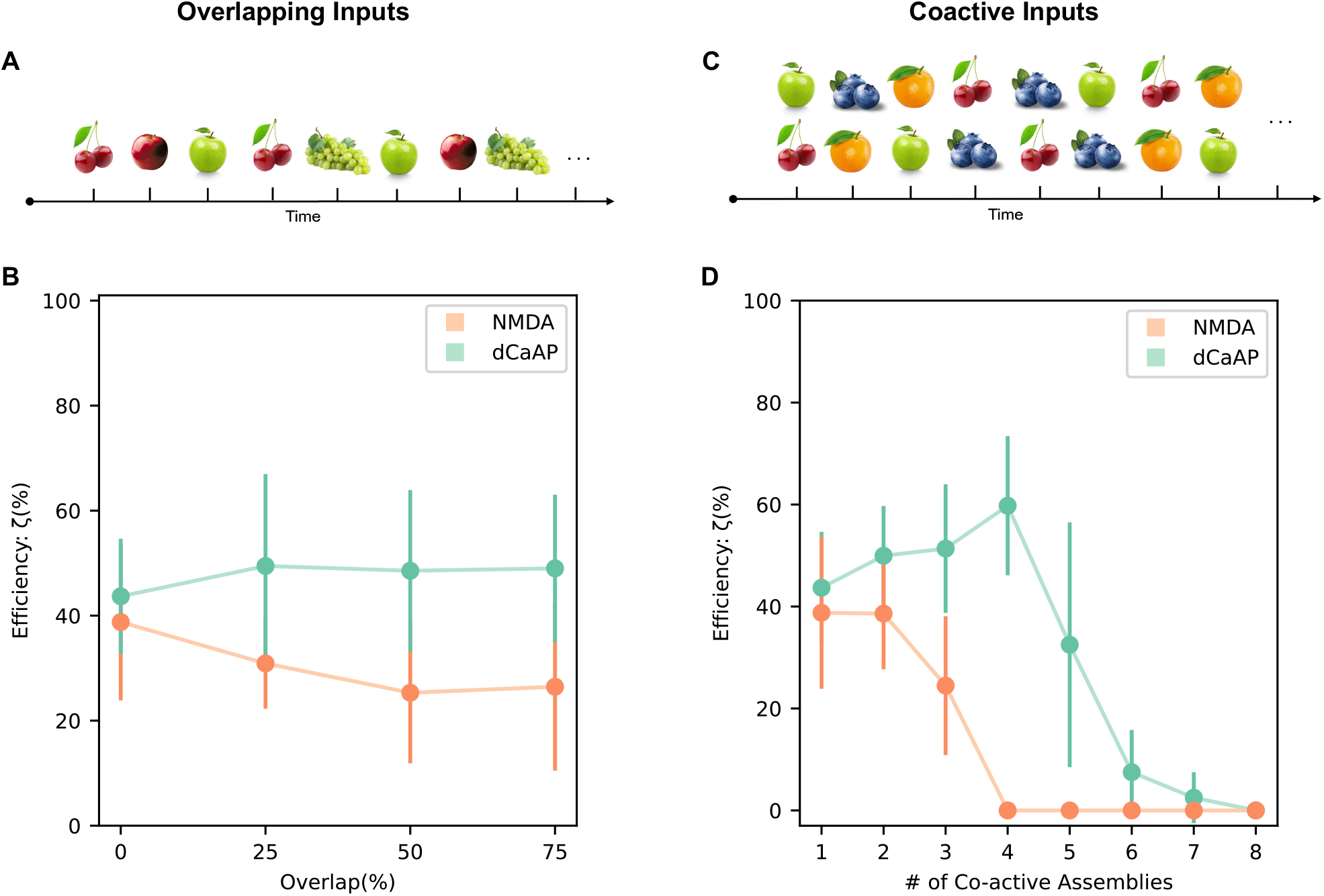
Overlapping and co-active inputs. *Overlapping Inputs:* **A)** The learning protocol is illustrated, where input assemblies share overlapping features such as color, shape, or fruit type. For example, the “green apple” assembly overlaps with “red apple” due to fruit type similarity and with “green grapes” due to shared color. **B)** As the overlap between assemblies increases, reflecting greater feature similarity, the NMDA model decreases in efficiency, while the dCaAP model maintains a higher efficiency. *Co-active Inputs:* **C)** The learning protocol involves presenting multiple fruits simultaneously, resulting in co-active input assemblies. **D)** Learning efficiency is compared between the NMDA and dCaAP models for different numbers of co-active assemblies. The NMDA model’s efficiency decreases sharply with the simultaneous presentation of more than one assembly, while the dCaAP model’s efficiency is higher for four co-active assemblies.

### Co-active inputs

In real-life scenarios, we are often exposed to multiple stimuli simultaneously. To simulate this, we examine the impact of presenting simultaneously active input assemblies, referred to as co-active assemblies, during the learning phase on the performance of the models. Unlike the first three protocols, in which only one assembly is active at each activation window, this protocol entails the random coactivation of multiple disjoint assemblies. For instance, in two co-active assemblies simulation, two randomly selected assemblies are active for 300 ms during the learning phase, while the remaining assemblies fire at a rate of 1 Hz. Following a 200-millisecond rest window, a new pair of assemblies is randomly chosen for simultaneous activation. A schematic representation of this protocol is shown in Figure 5C. This procedure is applied to co-active assemblies ranging from two to eight, with the results depicted in Figures 5D. Each data point in the figure represents the average efficiency value from 10 repetitions of the simulation. The results indicate that the NMDA model undergoes a rapid decline in efficiency with the addition of co-active assemblies, while the performance of the dCaAP model even improves with up to four co-active assemblies (see also Figures S5, S6 and S7 in the Supplementary Materials).

## Critical Factors for dCaAP Model Efficiency

In this section, we analyze the critical differences between the dCaAP and NMDA models that may underlie the superior performance of the dCaAP model in most investigated stimulation protocols, emphasizing their computational characteristics rather than their biological implications. The primary distinctions between the models include the triggering threshold, refractory period duration, dendritic spike amplitude, and response characteristics (all-or-none versus graded). To determine the key factors underlying the enhanced performance of the dCaAP model, we systematically changed these parameters and evaluated their impact. The results, presented in Figure 6, summarize the outcomes of 10 repetitions of the dCaAP model simulations for each parameter variation for different stimulation protocols. These include reducing the spike threshold from -36.0 mV to -55.0 mV (blue), shortening the refractory period by 75% (green), decreasing the dendritic spike amplitude by 20% (yellow), and disabling the graded amplitude response (pink). The analysis reveals that, across many protocols, an artificial reduction of either the dCaAP generation threshold or its maximal amplitude leads to a significant reduction in performance. This finding highlights the importance of dCaAP-specific properties in enabling a single neuron to learn and represent different input assemblies also for complex stimulation protocols.

**Figure 6:**
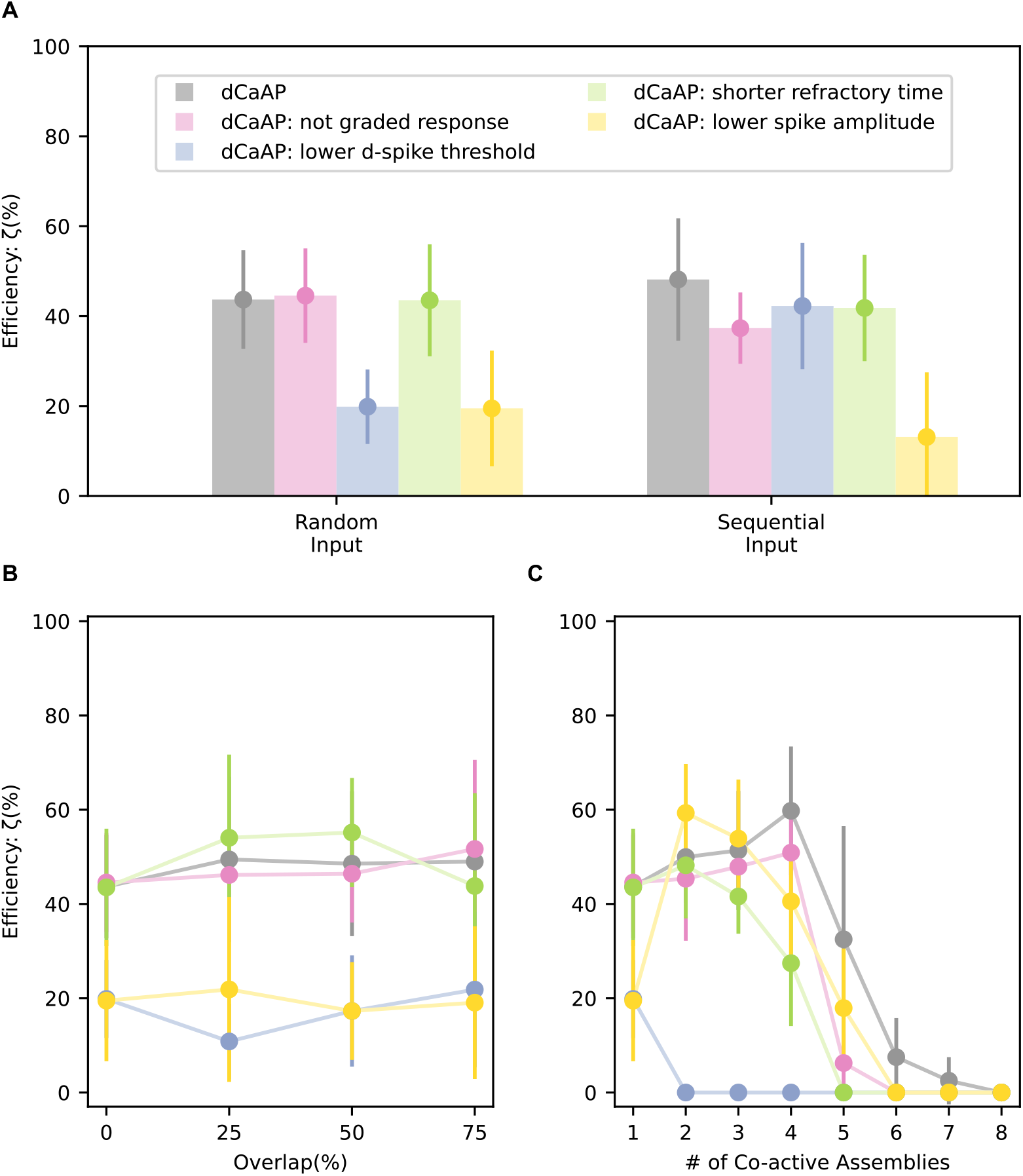
Key factors influencing the efficiency of the dCaAP model. To identify the key parameters responsible for the dCaAP model’s robust input disentanglement across different learning protocols, we altered four features: disabling the graded response (pink), lowering the dendritic spike threshold (blue), shortening the refractory period (green), and reducing the dendritic spike amplitude (yellow). The average results from 10 simulation repetitions for each variation are presented: **A)** random and sequential input protocols; **B)** overlapping inputs protocol; **C)** co-active input protocol.

## Discussion

The unique properties of dendritic calcium-mediated action potentials (dCaAPs)—such as their graded responses, high activation thresholds, and strong amplitudes [12]—led us to hypothesize that these spikes may play a critical role in synaptic clustering within dendrites. To test this hypothesis, we developed a computational model based on a previous study [28], which utilized NMDA spikes and synaptic plasticity to generate input clusters on dendritic branches; in contrast, our model incorporates the defining biophysical features of dCaAPs. We demonstrated that the interaction between the dCaAP and various synaptic plasticity rules—including structural, functional, and spike-timing-dependent plasticity (STDP)—promotes the formation of synaptic clusters on dendritic branches.

Our results demonstrate that the dCaAP-based model reliably forms stable synaptic clusters across different temporal input protocols. Specifically, both random and sequential presentations of input assemblies resulted in consistent and efficient branch-specific clustering. This characteristic could address a key limitation of conventional neural networks: catastrophic forgetting, in which the introduction of new information interferes with previously learned knowledge, leading to significant performance degradation or even complete erasure of prior information [21, 34, 36, 37, 40]. The brain exhibits exceptional continual learning capabilities, integrating new information without compromising prior knowledge [22, 40, 51, 53]. While hypotheses such as adaptive synaptic mechanisms and sleep-mediated unsupervised replay have been proposed to explain this phenomenon, the underlying neurobiological processes remain poorly understood [1, 51]. Despite the reasonable performance of the previous synaptic clustering model [28] in preventing the overwriting of previously acquired information, our findings reveal that under high input intensities, NMDA spikes fail to preserve learned data (see Figure S8 in the Supplementary Materials).

Remarkably, our model demonstrates enhanced robustness by employing dCaAPs. The enhanced efficacy of dCaAPs in facilitating synaptic clustering in our model is primarily attributed to their large spike amplitude, which promotes increased somatic spiking and enables more effective STDP-based learning on individual dendritic branches (Figure 6). This heightened branch-specific learning fosters competitive dynamics, ensuring that only a single branch forms strong connections with a given input assembly, while other branches remain available for future inputs, thereby reducing the likelihood of overwriting previously learned information. In contrast, the NMDA spike-based model, characterized by weaker amplitudes, generates fewer somatic spikes, leading to diminished competition among branches. Consequently, the learning capacity of the neuron is fully utilized, and the incorporation of new inputs necessitates the overwriting of existing knowledge. These results suggest that the large amplitude of dCaAPs is not merely a compensatory mechanism to mitigate dendritic attenuation, as proposed by Gidon et al. (2020) [12], but may also contribute significantly to preventing catastrophic forgetting.

Furthermore, experimental evidence shows that associative memories learned within short time intervals are encoded by overlapping populations of neurons [7, 17, 44, 45, 49]. Our results suggest that the dCaAP model can efficiently form synaptic clusters, even when assemblies overlap significantly, indicating that dendrites equipped with dCaAPs are capable of distinguishing between different elements of an associative memory.

In addition, a standout feature of our model is its remarkable ability to disentangle co-active neural assemblies, an essential function for navigating real-world environments where the brain continuously perceives inputs from various sources simultaneously. This capability is crucial for efficiently processing complex information. Our model achieves this by employing a high triggering threshold, ensuring that only temporally correlated “clusters” of input signals are strong enough to activate dendritic spikes. This approach promotes more focused and selective learning, enabling the model to better recognize meaningful patterns and associations. In contrast, the NMDA spike-based model, with its lower threshold, leads to more frequent and indiscriminate dendritic spikes, which can disrupt the fine-tuning of synaptic connections and hinder the process of synaptic clustering.

In summary, our results demonstrate that the dCaAPs support robust synaptic clustering, enhance learning stability, and prevent catastrophic forgetting, outperforming NMDA-based mechanisms, particularly under high input load and overlapping assemblies. These findings suggest that dCaAPs play a key role in enabling continual and selective learning. Future work could explore how dCaAP-like dynamics contribute to large-scale memory networks and their potential applications in neuromorphic computing or artificial intelligence.

## Author Contributions

CT, SSH, and SH conceived the research idea, wrote the manuscript, analyzed the simulation data and figures. SH developed the computational codes, performed simulations, and prepared the figures. Project supervision is done by CT and SSH.

