## Supplementary Materials for "Robust Input Disentanglement Through Dendritic Calcium-Mediated Action Potentials"

#### Efficiency factor

By the end of the learning phase, certain dendritic branches become specialized in recognizing distinct assemblies. This means that a branch forms strong connections predominantly with neurons from a single assembly. To compare the outcomes, we introduced an efficiency factor. Efficient learning is defined as the scenario where each dendritic branch of a neuron learns a distinct assembly, minimizing redundancy among branches. In other words, the number of dendritic branches connected to the same assembly should be as low as possible. Figure S1 illustrates the concept of the efficiency factor.

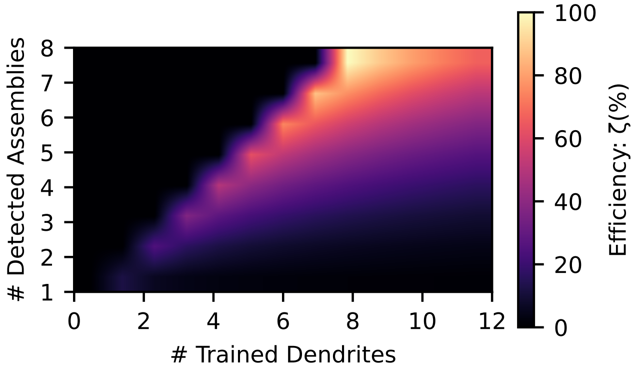


Figure S1: **Efficiency parameter.** A schematic representation of the defined efficiency factor is plotted as a function of the number of learned branches and detected assemblies. Since the number of detected assemblies cannot exceed the number of learned branches (as each branch can detect a maximum of one assembly if it is a 'learned branch'), the upper-diagonal region of the plot is shaded in black to indicate zero efficiency. Maximum efficiency is achieved when the model detects all 8 assemblies using the minimum required number of learned branches, namely 8. When the number of learned branches exceeds the number of detected assemblies, efficiency decreases, as this implies that an assembly is detected by multiple branches, indicating lower overall efficiency in the model.

### Simulations

Here, we describe the protocols for each simulation and present additional figures that support our findings discussed in the main text of the paper.

#### Random Inputs

In the first protocol, training was conducted over a period of 1000 seconds, equivalent to $10^6$ timesteps. During each training interval, a single assembly was activated at a high rate for 300 ms, followed by a 200 ms rest period when no assemblies were active. Subsequently, another assembly was randomly selected and activated at a high rate.

Figure S2 shows representative membrane potential traces from 4 dendritic branches of a dCaAP model during this training process. In the left column, plots represent the initial 10 seconds of training, while in the right column the final 10 seconds are displayed. The first row shows the activity of the 320 presynaptic neurons, while rows 2 through 5 present dendritic membrane potentials from four sample branches.

#### Overlapping Inputs

In this protocol, we trained the model with overlapping assemblies and assessed their performance. As shown in Figure S4, the dCaAP model can detect more assemblies, even when the assemblies are not completely disjoint and have a significant shared population.

#### Co-active Inputs

In this scenario, the activity of the assemblies is not temporally disjoint. This means that during each activity episode, multiple assemblies are simultaneously active at high frequencies. An example of two co-active assemblies is illustrated in Figure S6. During each learning window, two random assemblies are activated while the others are stimulated at a low frequency. For instance, the gray assembly is activated once with the brown assembly and subsequently with the green assembly. Figure S3 further illustrates the synaptic weights across these four branches, with each row representing a single branch. In the left column, each line denotes the cumulative weights of all connections from a specific assembly to the branch, while the right column depicts the total number of connections from each assembly to the branch. Over time, a single assembly progressively dominates each branch, as shown by an increase in both the number of connections and the overall synaptic weight. This pattern indicates that each branch is effectively learning and specializing in a distinct assembly.

The total synaptic weights and the number of connections to each assembly for this simulation are shown in Figure S7. The training results indicate that although two assemblies were activated in each training episode, each branch learned only one assembly. The results showing the number of detected assemblies for varying numbers of co-active assemblies are presented in Figure S5. These findings demonstrate that as the number of co-active assemblies increases, the performance of the dCaAP model improves to some extent (up to 4 co-active assemblies). In contrast, for the NMDA model, an increase in co-active assemblies leads to a sharp decline in the number of detected assemblies, significantly impacting the model's performance.

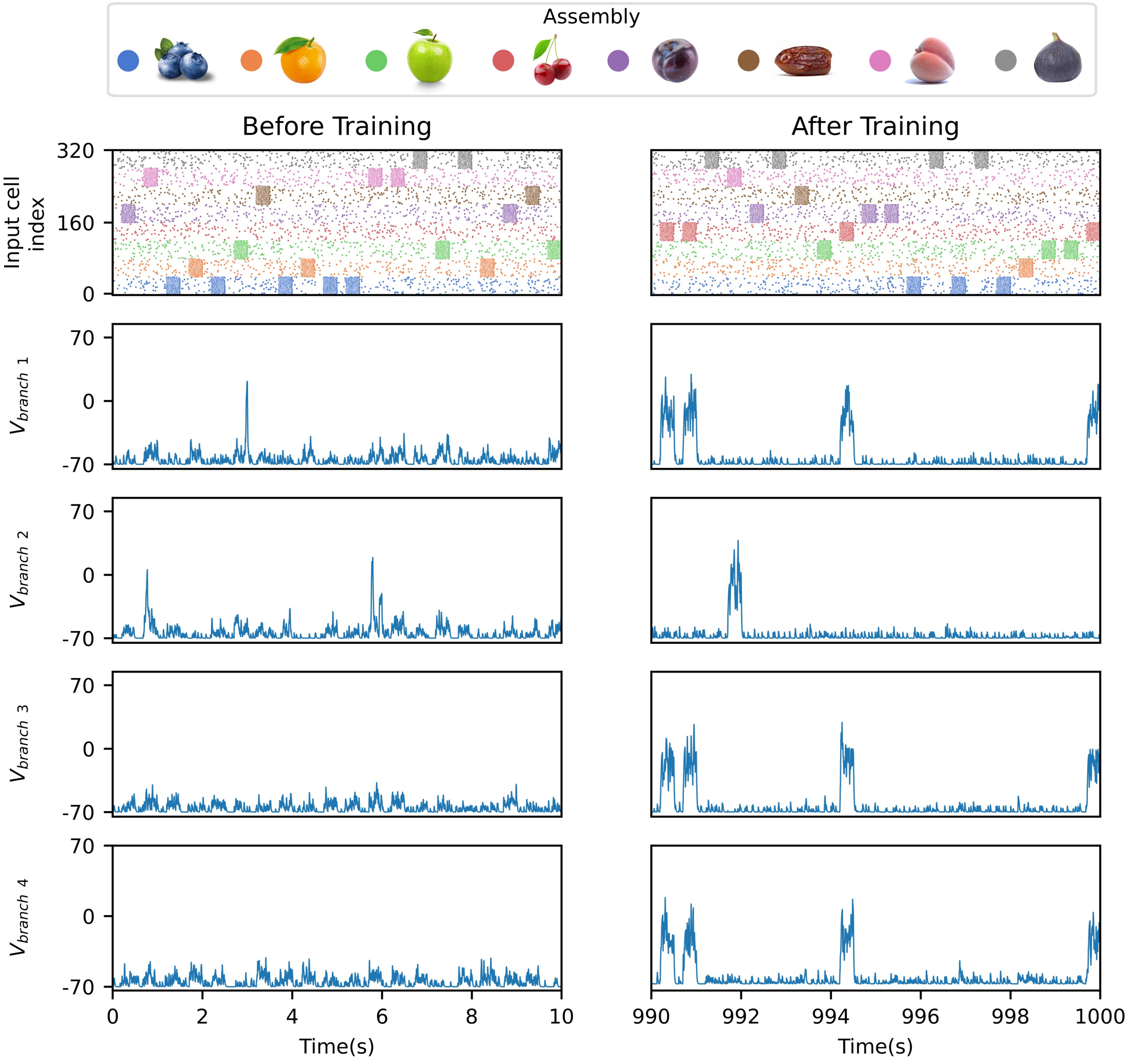


Figure S2: **Membrane potential traces during random inputs learning. Left:**This column illustrates the first 10 seconds of the training period. In the first row, the activity of 320 presynaptic neurons is represented, with each color denoting a different assembly. The membrane potentials of four sample branches are plotted in the subsequent rows. Since the branches have not yet learned the assemblies, their activities remain low at this stage. **Right:** This column displays the final 10 seconds of training, where the branches show heightened activity during the activation of specific assemblies. This behavior confirms that each branch has successfully learned an assembly.

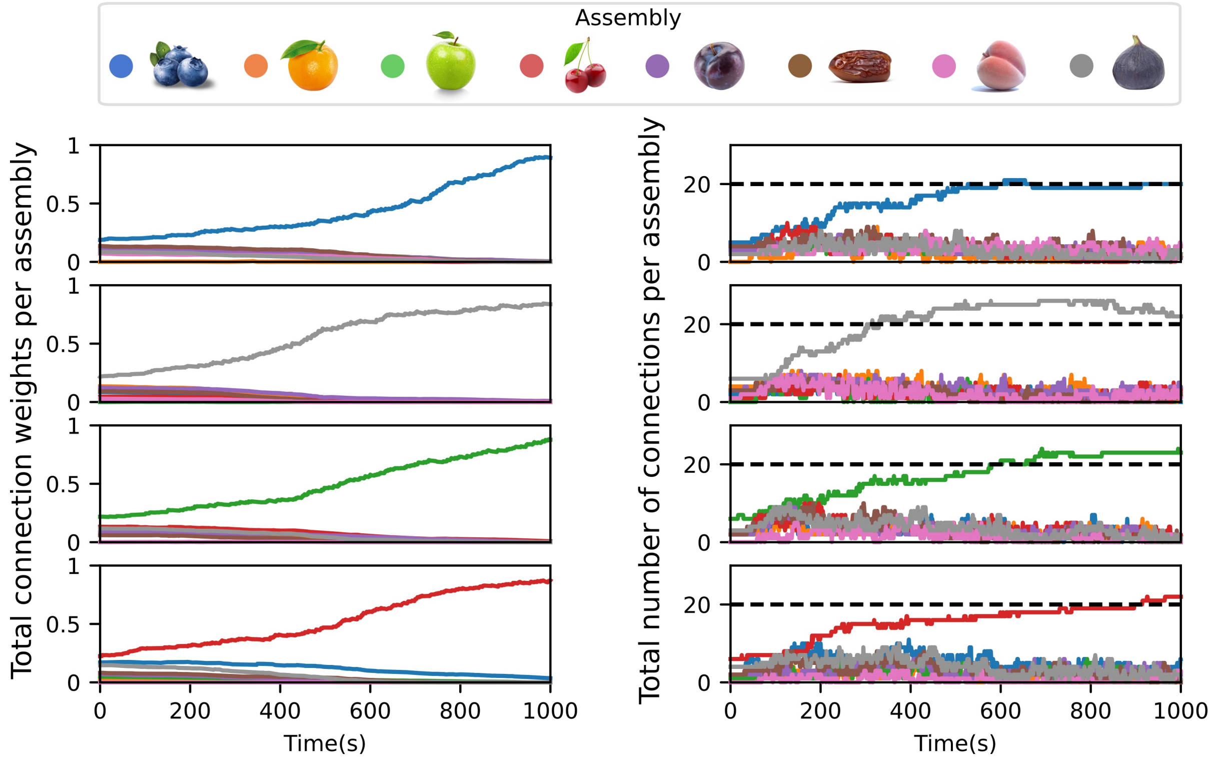


Figure S3: **Synaptic weights development during random inputs learning. Left:** Each row shows the sum of synaptic weights for all connections from a single assembly to a given branch. Over time, the weight sum for a single assembly on each branch increases, indicating its strengthening. The weights are scaled such that the maximum possible weight sum for any assembly is 1. **Right:** The total number of connections from each assembly to a single branch is shown over time. As observed, the connection count for the learned assembly reaches the structural limit (20 in this model) by the end of the training phase.

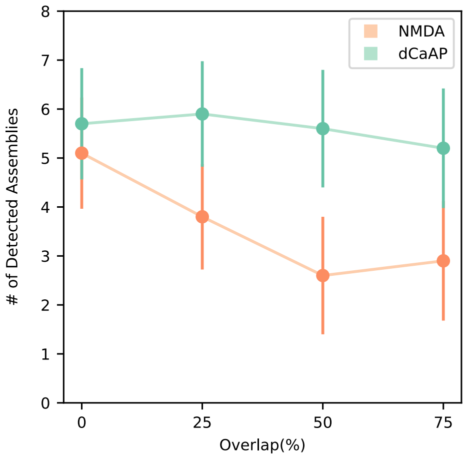


Figure S4: **Overlapping inputs.** The dCaAP model demonstrates superior performance.

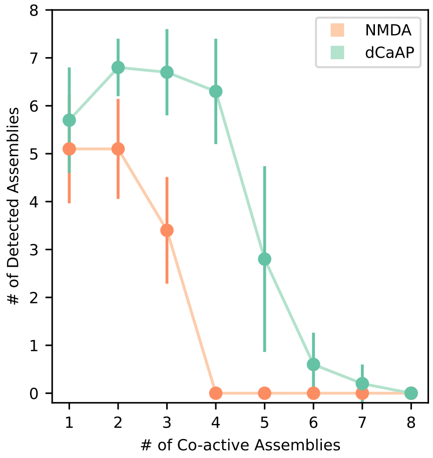


Figure S5: **Number of detected assemblies.** dCaAP model shows improved performance up to 4 co-active assemblies.

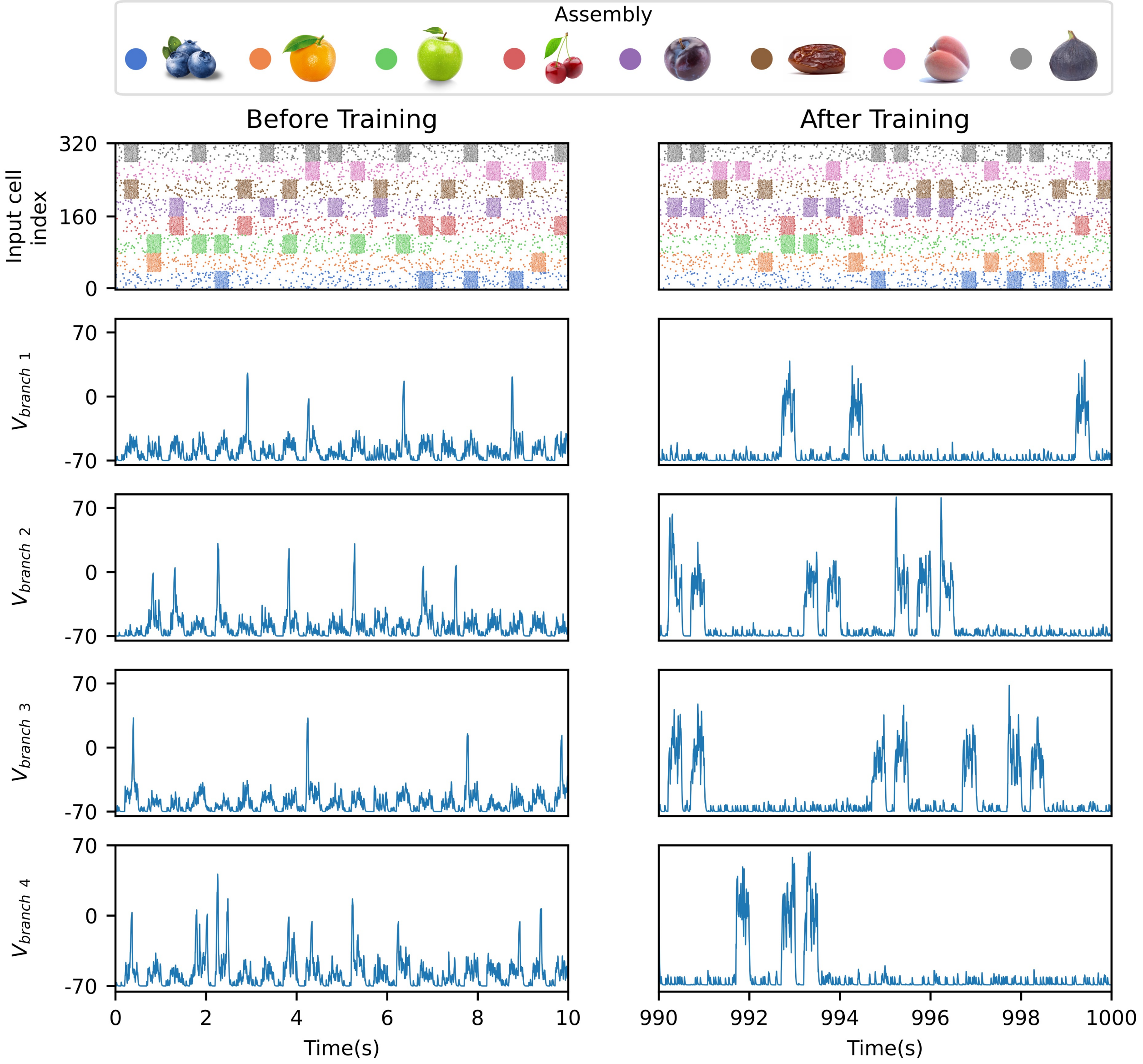


Figure S6: **Membrane potential traces during co-active inputs learning.** The first row represents the activity of the presynaptic cells, while the plots in each subsequent row depict the membrane potential of a dendritic branch. **Left:** The first 10 seconds of learning with two co-active assemblies show unspecialized branch activity during the presentation of distinct assemblies. **Right:** The final 10 seconds of learning demonstrate that each branch has successfully learned one assembly. This is evident from the branch's activity during the presentation of the corresponding assembly.

### Impact of Input Current Intensity

An influential parameter in the clustering efficiency of both models is current intensity. Figure S8 illustrates the efficiency trends of both models as current intensity increase in the random

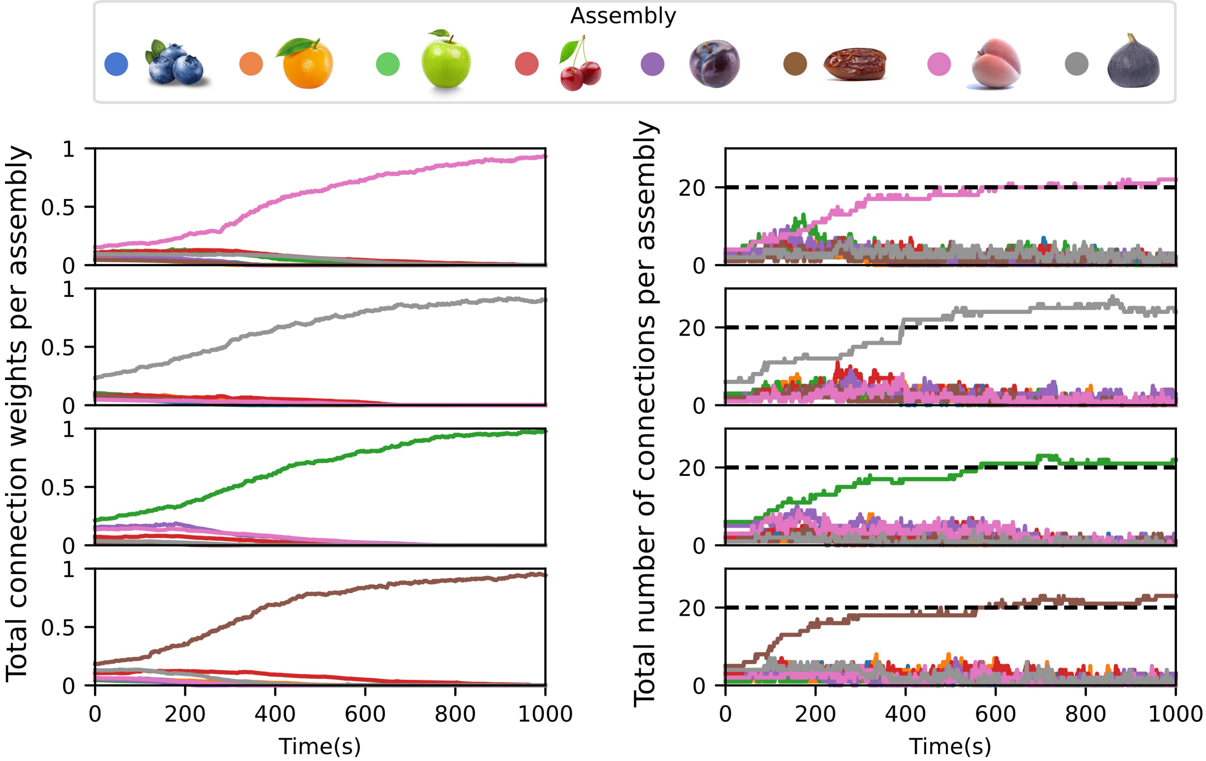


Figure S7: **Synaptic weights development during co-active inputs learning. Left:** The sum of the weights of all connections to each assembly on each branch is presented. All displayed branches have learned a distinct assembly. **Right:** By the end of the simulations, the total number of connections to the learned assembly on each branch reaches the structural limit (here is 20 connections).

input training protocol. The results indicate that the efficiency of the NMDA model declines with increasing intensity. Conversely, at lower intensities, the dCaAP model exhibits reduced efficiency compared to the NMDA model; however, at higher intensities, the dCaAP model surpasses the NMDA model in performance. This behavior can be attributed to the higher activation threshold of the dCaAP model. At lower intensities, the elevated threshold of the dCaAP model makes it more challenging to trigger dendritic spikes, resulting in limited learning and allowing the NMDA model to outperform it. Conversely, the lower threshold of the NMDA model causes excessive activity during high-intensity paradigms, which reduces its learning efficacy. In contrast, the higher threshold of the dCaAP model promotes more selective spike activation, leading to improved performance under higher intensity conditions.

These findings are consistent with experimental work by Magó et al. (2021) [1], who identified a calcium dendritic spike in CA3 pyramidal cells of rats with properties akin to those of dCaAP. Their study demonstrated that these spikes occur more frequently as input current intensity increases. Our computational results suggest that these spikes enhance dendritic input clustering performance under similar high-intensity input conditions. Thus, the increased prevalence of dCaAP dendritic spikes may represent a compensatory mechanism that enables dendrites to sustain high clustering efficiency at elevated intensities.

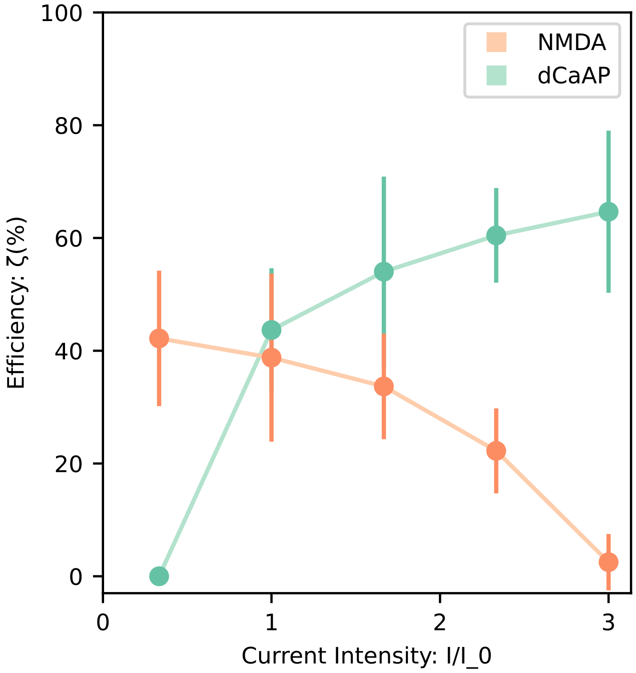


Figure S8: **Effect of input current intensity.** At lower intensities, the NMDA model performs better, whereas at higher intensities, the dCaAP model demonstrates greater efficiency.
